# A cloning-free method for CRISPR/Cas9-mediated genome editing in fission yeast

**DOI:** 10.1101/239673

**Authors:** Xiao-Ran Zhang, Jia-Bei He, Yi-Zheng Wang, Li-Lin Du

## Abstract

The CRISPR/Cas9 system, which relies on RNA-guided DNA cleavage to induce site-specific DNA double-strand breaks, is a powerful tool for genome editing. This system has been successfully adapted for the fission yeast *Schizosaccharomyces pombe* by expressing Cas9 and the single-guide RNA (sgRNA) from a plasmid. In the procedures published to date, the cloning step that introduces a specific sgRNA target sequence into the plasmid is the most tedious and time-consuming. To increase the efficiency of applying the CRISPR/Cas9 system in fission yeast, we here developed a cloning-free procedure that uses gap repair in fission yeast cells to assemble two linear DNA fragments, a gapped Cas9-encoding plasmid and a PCR-amplified sgRNA insert, into a circular plasmid. Both fragments contain only a portion of the *ura4* or *bsdMX* marker so that only the correctly assembled plasmid can confer uracil prototrophy or blasticidin resistance. We show that this gap-repair-based and cloning-free CRISPR/Cas9 procedure permits rapid and efficient point mutation knock-in, endogenous N-terminal tagging, and genomic sequence deletion in fission yeast.

## INTRODUCTION

In recent years, the CRISPR/Cas9 system has been successfully utilized for efficient genome editing in various eukaryotic organisms (Sander and Joung 2014; Doudna and Charpentier 2014; Wang et al. 2016). This system usually contains two components: one is the Cas9 protein; the other is a single-guide RNA (sgRNA), which is composed of a scaffold sequence for Cas9-binding and a user-defined 20-nucleotide target sequence. Together, Cas9 and sgRNA form an RNA-guided endonuclease that creates a double-strand break (DSB) in the target DNA 3 bp upstream of the protospacer-adjacent motif (PAM), which is NGG for the most commonly used Cas9 protein from *Streptococcus pyogenes* (Jinek et al. 2012). During the repair of the Cas9-generated DSBs, the desired genome editing outcomes can be achieved through either random mutations introduced by the error-prone non-homologous end joining (NHEJ) pathway or precise mutations introduced by the homologous recombination (HR) pathway (Sander and Joung 2014).

In deploying the CRISPR/Cas9 system, the expression of the sgRNA poses a special challenge, because the most commonly used promoters, which are RNA polymerase II-transcribed, do not allow the production of sgRNAs with precise 5′ end and 3′ end. For this reason, sgRNAs have usually been expressed from RNA polymerase III-transcribed promoters (Mali et al. 2013). However, RNA polymerase III-transcribed promoters are not well characterized in many organisms and often require particular nucleotides at the 5′ end of sgRNAs (Gao and Zhao 2014; Sander and Joung 2014). In fission yeast, this challenge has been solved elegantly by the combined use of the promoter/leader sequence of a RNA polymerase II-transcribed RNA-encoding gene, *rrk1*, and a self-cleavage hammerhead ribozyme (Jacobs et al. 2014).

In the published procedures of applying the CRISPR/Cas9 technology in fission yeast, a plasmid-cloning step is always undertaken each time a new sgRNA is needed (Jacobs et al. 2014; Rodriguez-Lopez et al. 2016; Fernandez and Berro 2016). This step is a multi-day endeavor that includes manipulations of DNA, transforming *E. coli* cells, plasmid preparation, and plasmid verification by sequencing. Because the cloned plasmid is destined to be delivered into the fission yeast cells and likely serves no further use once the intended genome editing is achieved, we envisioned that the cloning step can be omitted by assembling the Cas9-sgRNA plasmid through the highly efficient in vivo gap repair process in the fission yeast cells (Kostrub et al. 1998; Kelly and Hoffman 2002; Colon and Walworth 2004; Matsuo et al. 2010; Chino et al. 2010). In this work, we designed and implemented a gap-repair-based and cloning-free procedure for applying the CRISPR/Cas9 technology in fission yeast. We validated its use in point mutation knock-in, endogenous N-terminal tagging, and genomic sequence deletion.

## MATERIALS AND METHODS

### Fission yeast strains and culturing conditions

The fission yeast strain LD260 (*h^−^ ura4-D18 leu1-32 his3-D1*) was from our laboratory strain collection, and the naturally isolated strain JB938 was acquired from the Yeast Genetic Resource Center of Japan (YGRC/NBRP) (http://yeast.nig.ac.jp/) (Jeffares et al. 2015). Cells were cultured in pombe minimal medium with glutamate (PMG), or a yeast extract-based rich medium (YES) at 30°C or other temperatures as noted. The compositions of these media and standard genetic methods were as described (Forsburg and Rhind 2006).

### Plasmid construction

Plasmids used in this study are listed in Table 1, and primers used for plasmid construction are listed in Table S1.

**Table 1.** Plasmids used in this study

| Plasmid | Description | Addgene ID |
| --- | --- | --- |
| pMZ374 | A Cas9-encoding plasmid that contains an intact *ura4* marker, and is used in cloning-based procedures (Jacobs et al. 2014). | 59896 |
| pDB4280* | A Cas9-encoding plasmid containing the 5' portion of the *ura4* marker and the *rrk1* promoter/leader. When linearized by NotI, the resultant 10,201-bp DNA serves as the gapped plasmid in the split-*ura4* system. | 98699 |
| pDB4282* | A plasmid containing the 3' portion of the *ura4* marker and the sgRNA elements downstream of the target sequence. When digested by NotI, a 1,354-bp fragment serves as the PCR template for generating the sgRNA insert in the split-*ura4* system. | 98701 |
| pDB4279* | A pMZ374-derived plasmid with the *ura4* marker changed to the *bsdMX* marker. | 98698 |
| pDB4281* | A Cas9-encoding plasmid containing the 5' portion of the *bsdMX* marker and the *rrk1* promoter/leader. When linearized by NotI, the resultant 9,800-bp DNA serves as the gapped plasmid in the split-*bsdMX* system. | 98700 |
| pDB4283* | A plasmid containing the 3' portion of the *bsdMX* marker and the sgRNA elements downstream of the target sequence. When digested by NotI, a 931-bp fragment serves as the PCR template for generating the sgRNA insert in the split-*bsdMX* system. | 98702 |
| pDB4284* | An *rpl42*-targeting Cas9-sgRNA plasmid generated by cloning. |  |

The plasmids constructed in this study are indicated by an asterisk.

We constructed two plasmids for the split-*ura4* system. For the first plasmid (pDB4280), we modified the plasmid pMZ374 (Jacobs et al. 2014), by replacing a 1,191-bp sequence between the *rrk1* promoter/leader and the 160th codon of *ura4* gene with a NotI site using the Quikgene method with oligos P127 and P128 (Mao et al. 2011). For the second plasmid (pDB4282), a 1,324-bp sequence containing the 3′ part of the *ura4* marker starting from the 108th codon, the hammerhead ribozyme coding sequence, and the sgRNA scaffold sequence, was amplified from pMZ374 using oligos P75 and P82, and then cloned into the pEASY-blunt vector (Transgene, China).

For the split-*bsdMX* system, we first switched the *ura4* marker in pMZ374 to a drug resistance marker *bsdMX* by gap repair in fission yeast (Fennessy et al. 2014). The *bsdMX* marker, which confers blasticidin S resistance, was amplified from an MDR-supML strain provided by Shigehiro Kawashima (Aoi et al. 2014), using oligos P123 and P124, which contain sequences that base pair with sequences flanking the *ura4* marker in pMZ374 (38-bp upstream sequence and 36-bp downstream sequence). The *bsdMX* PCR product was co-transformed with StuI digested pMZ374 into fission yeast cells, and the episomal plasmid rescued from a resultant blasticidin-resistant colony was named pDB4279. A 770-bp sequence between the *rrk1* promoter/leader and the 89th codon of the *bsdMX* gene in pDB4279 was replaced with a NotI site using the Quikgene method with oligos P129 and P130. The resultant plasmid was named pDB4281. A 901-bp fragment containing the 3′ part of the *bsdMX* marker starting from the 38th codon, the hammerhead ribozyme coding sequence, and the sgRNA scaffold sequence, was amplified from pDB4279 with oligos P131 and P82, and then cloned into pEASY-blunt vector (Transgene, China), generating the pDB4283 plasmid.

### Preparation of the gapped plasmid

For the split-*ura4* system, a gapped plasmid (10,201 bp) containing the Cas9-coding sequence and the 5′ part of the *ura4* marker was generated by linearizing pDB4280 with NotI digestion.

For the split-*bsdMX* system, a gapped plasmid (9,800 bp) that contains the Cas9-coding sequence and the 5′ part of the *bsdMX* marker was generated by linearizing pDB4281 with NotI digestion.

### Preparation of the PCR templates for sgRNA insert amplification

For the split-*ura4* system, the PCR template (1,354 bp) for amplifying sgRNA inserts was generated from pDB4282 by NotI digestion. The template contains the 3′ part of the *ura4* marker, the hammerhead ribozyme sequence, and the sgRNA scaffold sequence, but not the sgRNA target sequence and the *rrk1* promoter/leader sequence.

For the split-*bsdMX* system, the PCR template (931 bp) for amplifying sgRNA inserts was generated from pDB4283 by NotI digestion. The template contains the 3′ part of the *bsdMX* marker, the hammerhead ribozyme sequence, and the sgRNA scaffold sequence, but not the sgRNA target sequence and the *rrk1* promoter/leader sequence.

### Target sequence selection

The sgRNA target sequence is the 20 nucleotides upstream of the PAM sequence, which is NGG, where N is any nucleotide (Jinek et al. 2012). When more than one NGG sequence can suit the need of genome editing, sgRNA target sequences were selected based on their “on-target scores” calculated by an sgRNA designing software at benchling.com (Doench et al. 2016). In our experience, a high “on-target score” did not always translate into a high editing efficiency. Thus, if possible, we usually chose two different sgRNA target sequences for each editing task. The sgRNA target sequences used in this study are listed in Table 2.

**Table 2.**
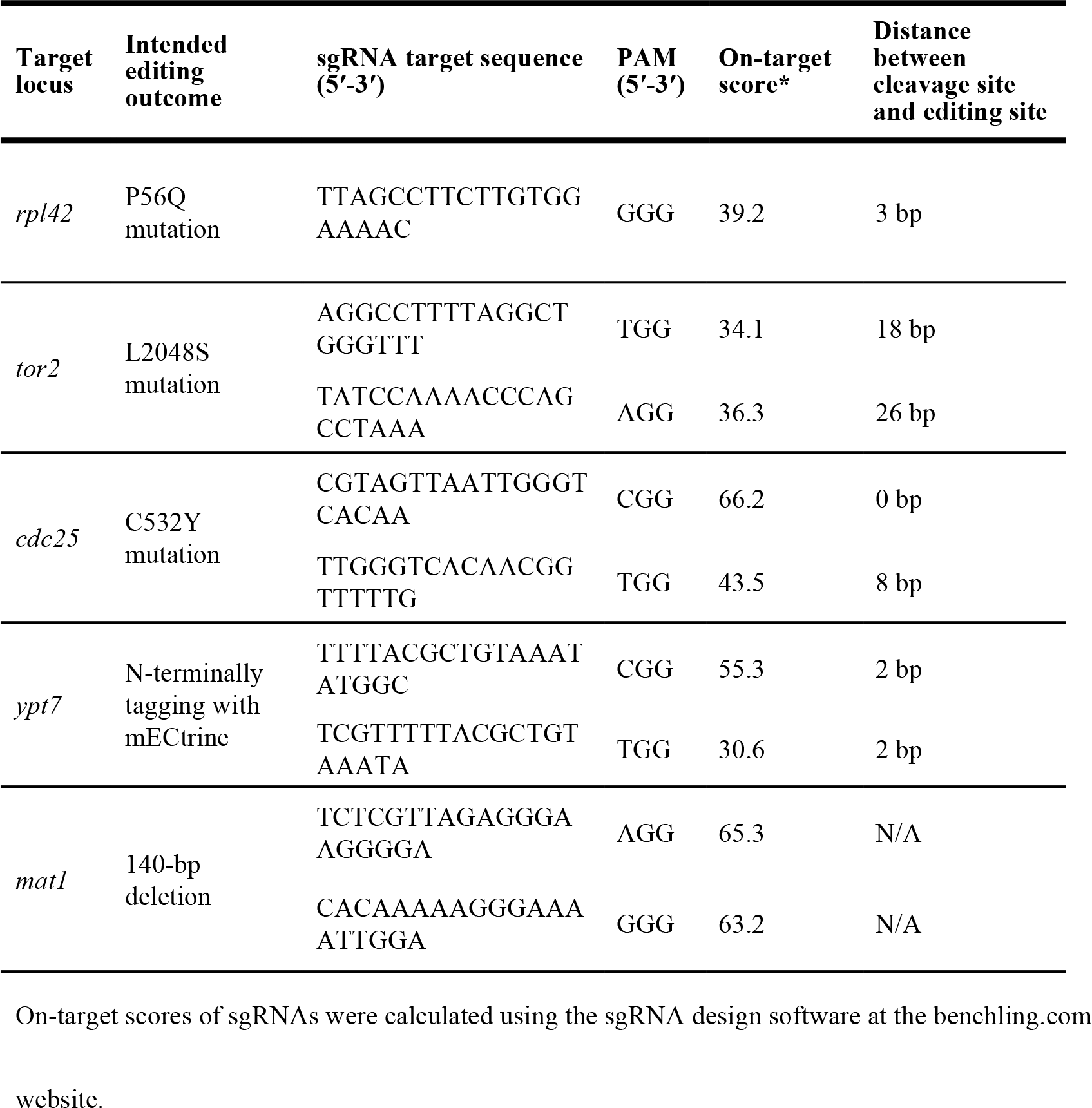
Genome editing performed in this study

### sgRNA insert amplification

The sgRNA target sequence (5′-3′) was incorporated into a 83-nt sgRNA primer (5’-ATAGTTGCTGTTGCCAAAAAACATAACCTGTACCGAAGAA*NNWNNNNNNNNNNNNNNNN*gttttagagctagaaatagcaag-3’), which contains a 40-nt sequence (in capital letters) from the *rrk1* leader, followed by the 20-nt sgRNA target sequence (shown as 20 Ns in italic), and a 23-nt sequence from the sgRNA scaffold sequence (in lower case letters).

Using a high-fidelity DNA polymerase, a common primer (P75, 5′-CATCTGGTGTGTACAAAATTG-3’, for the split-*ura4* system, or P131, 5′-GGCCGCATCTTCACTGGTGTC-3′, for the split-*bsdMX* system) was paired with the sgRNA primer to amplify the sgRNA insert from the PCR template aforementioned. To facilitate gap repair, the sgRNA insert ends with sequences that overlap with the terminal sequences of the gapped plasmid. For the split-*ura4* system, the sgRNA insert (1,375 bp) contains a 158-bp overlapping sequence (in the *ura4* selection marker) at one end and a 40-bp overlapping sequence (in the *rrk1* leader) at the other end. For the split-*bsdMX* system, the sgRNA insert (952 bp) contains a 156-bp overlapping sequence (in the *bsdMX* selection marker) at one end and a 40-bp overlapping sequence (in the *rrk1* leader) at the other end.

### Design of donor DNA

To achieve a precise genome-editing outcome, we used donor DNA carrying the intended mutation and if the intended mutation does not abolish Cas9 cleavage, at least one additional synonymous cleavage-blocking mutation locating in the last 10 nucleotides of the sgRNA target sequence or in the PAM sequence. For knock-in or knock-out, the donor DNA was provided as a pair of 90-nt synthetic oligos whose sequences are reverse complementary to each other. For knock-in, the desired mutation(s) were placed approximately in the middle of the 90-nt sequence. For knock-out, two 45-bp sequences flanking the region to be deleted were joined together to serve as the donor. For N-terminal tagging, the donor DNA was provided as a PCR product, which contains homology arm sequences flanking the tag sequence. The donor oligos and primers used in this study are listed in Table S2.

### Transformation, selection, and phenotype assessment

The gapped plasmid, the sgRNA insert PCR product, and the donor DNA (two 90-nt synthetic oligos or a PCR product) were co-transformed into an appropriate fission yeast strain using the lithium acetate/single-stranded carrier DNA/polyethylene glycol method (Gietz and Woods 2006). The split-*ura4* system requires the host strain to harbor the *ura4-D18* allele, which is a complete deletion of the sequence in the *ura4* marker (Grimm et al. 1988). The split-*bsdMX* system requires the host strain to not already have a blasticidin S resistance marker. 5 OD600 units of cells were used in each transformation. A donor-free transformation was always included as a control to assess the effect of the donor on the transformation efficiency. A strong donor-dependent increase of transformation efficiency usually correlated with high genome editing efficiency. An optional but useful control is a transformation of a plasmid that contains a full-length selection marker but not an sgRNA target sequence (pMZ374 for the split-*ura4* system and pDB4279 for the split-*bsdMX* system). Such a transformation is expected to yield a high transformation efficiency due to the lack of Cas9 cleavage of the genome. Ineffective sgRNA can result in a similarly high transformation efficiency even without the donor DNA. Another optional control that serves a similar purpose is the co-transformation of the gapped plasmid with an sgRNA insert amplified using a control sgRNA primer. We used 5′-ATAGTTGCTGTTGCCAAAAAACATAACCTGTACCGAAGAA*tgggcttaactcaattcttgtgggttatctctct*gttttagagctagaaatagcaag-3′ as the control sgRNA primer to amplify an sgRNA insert, which through gap repair generates the same plasmid as pMZ374 or pDB4279.

For the split-*ura4* system, uracil prototrophic transformants were selected on PMG plates with necessary supplements but lacking uracil. Colonies usually formed after incubating at 30°C for 6-8 days. This longer than usual time taken for transformant colonies to form is due to the growth inhibitory effect of Cas9 on fission yeast cells (Jacobs et al. 2014).

For the split-*bsdMX* system, transformants were selected on YES plates supplemented with 30 μg/ml of blasticidin S hydrochloride (Wako Pure Chemical Industries, Ltd., catalog number 029-18701). To enhance the transformation efficiency, instead of the common practice of plating cells onto YES prior to replica-plating onto selective media for antibiotic marker selection, we performed a liquid phase recovery in MSL-N liquid medium for 24 hours prior to plating on blasticidin-containing plates (Fennessy et al. 2014). Colonies usually formed after incubating at 30°C for 4-6 days.

The correct genome editing outcomes were scored by assessing the expected phenotypes of the intended mutations. For the *rpl42-P56Q* mutation, the expected resistance to cycloheximide (CYH) was assessed by replica plating onto YES plates containing 100 μg/ml of CYH; For *tor2-L2048S* and *cdc25-C532Y* mutations, the expected temperature-sensitive phenotype was analyzed by replica-plating onto two YES plates and then incubating one of them at the permissive temperature (25°C) and the other at the restrictive temperature (36°C). For N-terminal tagging of Ypt7 with a fluorescent protein, the clones were grown in liquid PMG medium containing necessary supplements and then observed with microscope. For converting a homothallic *h90* strain to heterothallic strains by introducing the *mat1*-Δ*17* mutation (Arcangioli and Klar 1991), the transformants were replica plated onto SPAS plates with necessary supplements to induce mating, meiosis, and sporulation, and then stained with iodine vapor, which stains spores dark brown (Forsburg and Rhind 2006).

As has been observed by others (Jacobs et al. 2014; Rodriguez-Lopez et al. 2016), the rare fast-growing transformant colonies, which appeared to have escaped from the growth inhibitory effect of Cas9, usually do not harbor the intended genome-editing outcome, probably because Cas9 is absent or not functional in these clones. Such colonies were avoided when picking random colonies for phenotype assessment (e.g. in the case of Ypt7 tagging), and were not counted when quantitating transformation and genome editing efficiencies.

Because of the growth inhibitory effect of Cas9 on fission yeast cells, the removal of the episomal Cas9-sgRNA plasmid can be quickly achieved by growing the transformant clones on a non-selective medium.

### Light microscopy

Live cell imaging was performed using a DeltaVision PersonalDV system (Applied Precision) equipped with a CFP/YFP/mCherry filter set (Chroma 89006 set) and a Photometrics CoolSNAP HQ2 camera. Images were acquired with a 100x 1.4-NA objective lens and were analyzed with the softWoRx program.

### Reagent availability

Five plasmids created in this study (pDB4279, pDB4280, pDB4281, pDB4282, and pDB4283) have been deposited at Addgene (plasmid number 98698, 98699, 98700, 98701, and 98702, respectively). Plasmid map files in GenBank format are provided as Supplementary Data Files 1-5.

### RESULTS

#### A gap-repair-based CRISPR/Cas9 procedure can efficiently knock in a point mutation in fission yeast

To make the CRISPR/Cas9 system more efficient in fission yeast, we developed a cloning-free procedure, which assembles the Cas9-sgRNA plasmid by gap repair in fission yeast cells (Fig. 1). We call one version of this procedure the split-*ura4* system, which utilizes two plasmids pDB4280 and pDB4282 (Table 1), both derived from the *ura4*-marked plasmid pMZ374 (Jacobs et al. 2014). Digesting these two plasmids with NotI generates the gapped plasmid and the PCR template for sgRNA insert amplification, respectively. Using a common primer and an 83-nt sgRNA-specific primer to perform PCR, an sgRNA insert containing a specific target sequence is generated by PCR amplification. At the two ends of an sgRNA insert are sequences overlapping with the gapped plasmid. Upon co-transformation of the gapped plasmid and the sgRNA insert into a *ura4-D18* fission yeast strain, a plasmid expressing both Cas9 and sgRNA is assembled in vivo by gap repair. Importantly, neither the gapped plasmid nor the sgRNA insert has a complete *ura4* marker, so that only the correctly assembled plasmid can result in the transformation of a uracil auxotrophic *ura4-D18* strain to a uracil prototrophic one. If provided with an appropriate donor DNA for homologous recombination-mediated repair of the Cas9-induced DSB, this cloning-free CRISPR/Cas9 procedure should permit rapid and precise genome editing.

**Figure 1.**
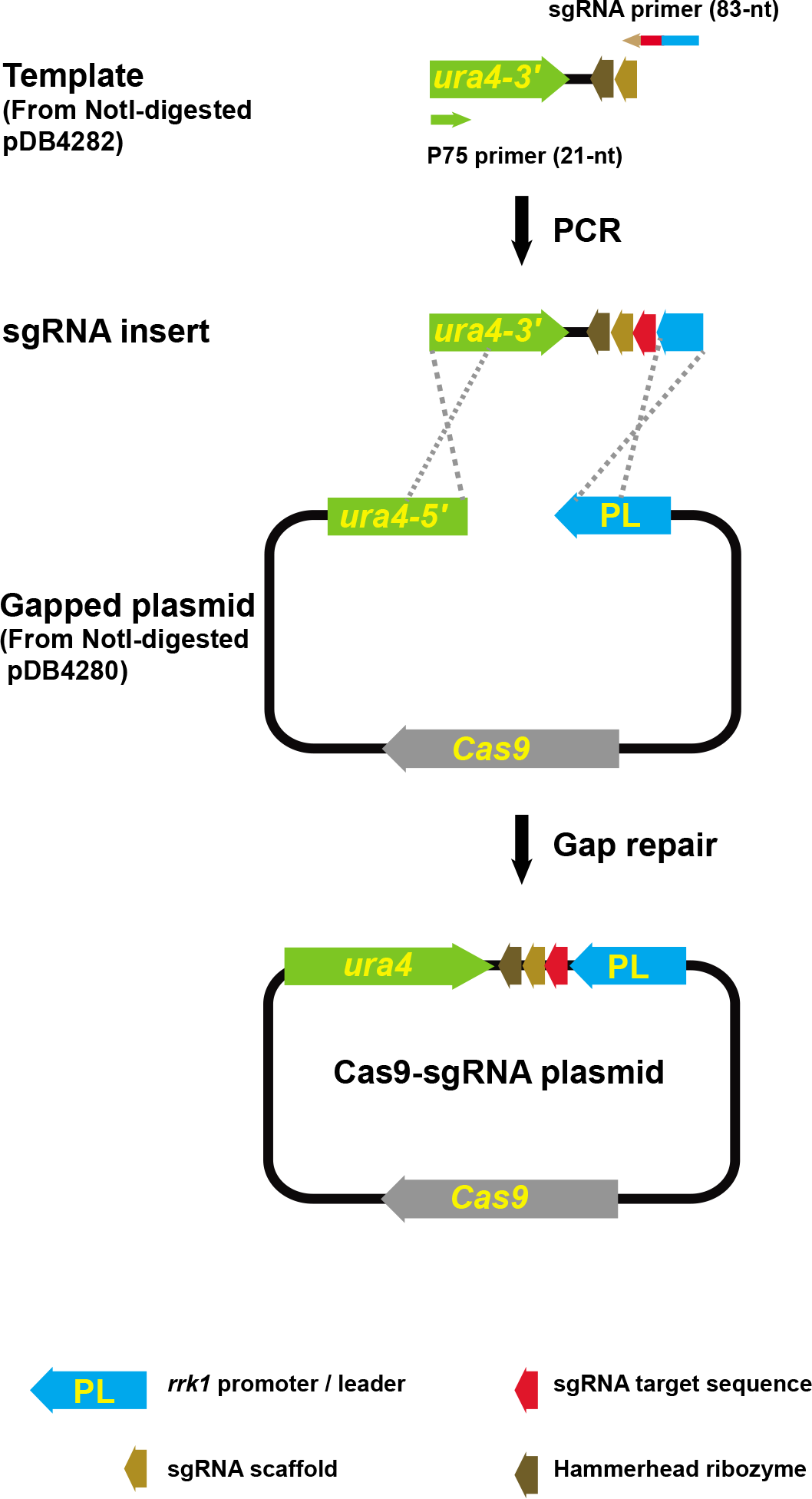
The cloning-free procedure that generates the Cas9-sgRNA plasmid through gap repair. In this procedure, the sgRNA target sequence, depicted in red, is synthesized as part of an 83-nt oligo (sgRNA primer), then incorporated into a linear DNA fragment (sgRNA insert) by PCR, and eventually recombined into a circular plasmid by gap repair in fission yeast cells. The overlapping sequences involved in gap repair are indicated by the crossed dashed lines. To prevent gap-repair-independent transformation, the gapped plasmid and the sgRNA insert contain only the 5′ and the 3′ parts of the selection marker, respectively. Only proper gap repair can reconstitute the complete marker (*ura4* in the split-*ura4* system and *bsdMX* in the split-*bsdMX* system). The split-*ura4* system is depicted here (not drawn to scale).

To test this procedure, we used it to knock-in the *rpl42-P56Q* mutation, which is known to confer cycloheximide resistance (CYH^R^) (Shirai et al. 2010; Roguev et al. 2007). The nucleotides to be mutated (a CCC codon for proline) coincide with a PAM sequence (GGG on the opposite strand), and thus we were able to use an sgRNA that induces Cas9 cleavage only 3 bp away from the editing site (Fig. 2A and Table 2). Ura^+^ transformant colonies that contain the knock-in mutation were identified by replica plating transformant colonies onto YES plates containing CYH (Fig. 2B). The ratio of the number of CYH^R^ colonies to the number of Ura+ transformant colonies (CYH^R^/Ura^+^) was taken as the knock-in or editing efficiency. Using a pair of 90-nt complementary oligos as donor DNA, the gap repair procedure achieved a high editing efficiency (84%), comparable to that obtained using a cloned Cas9-sgRNA plasmid (Fig. 2B and 2C). The presence of donor DNA was not only critical for the knock-in to occur, but also notably increased the transformation efficiency, presumably because the knock-in mutation blocks Cas9-mediated DNA cleavage, which is detrimental to the cells. Unexpectedly, the gap repair procedure resulted in more than 10-fold higher transformation efficiencies than the procedure using the cloned plasmid. The exact reason behind this phenomenon is unclear.

**Figure 2.**
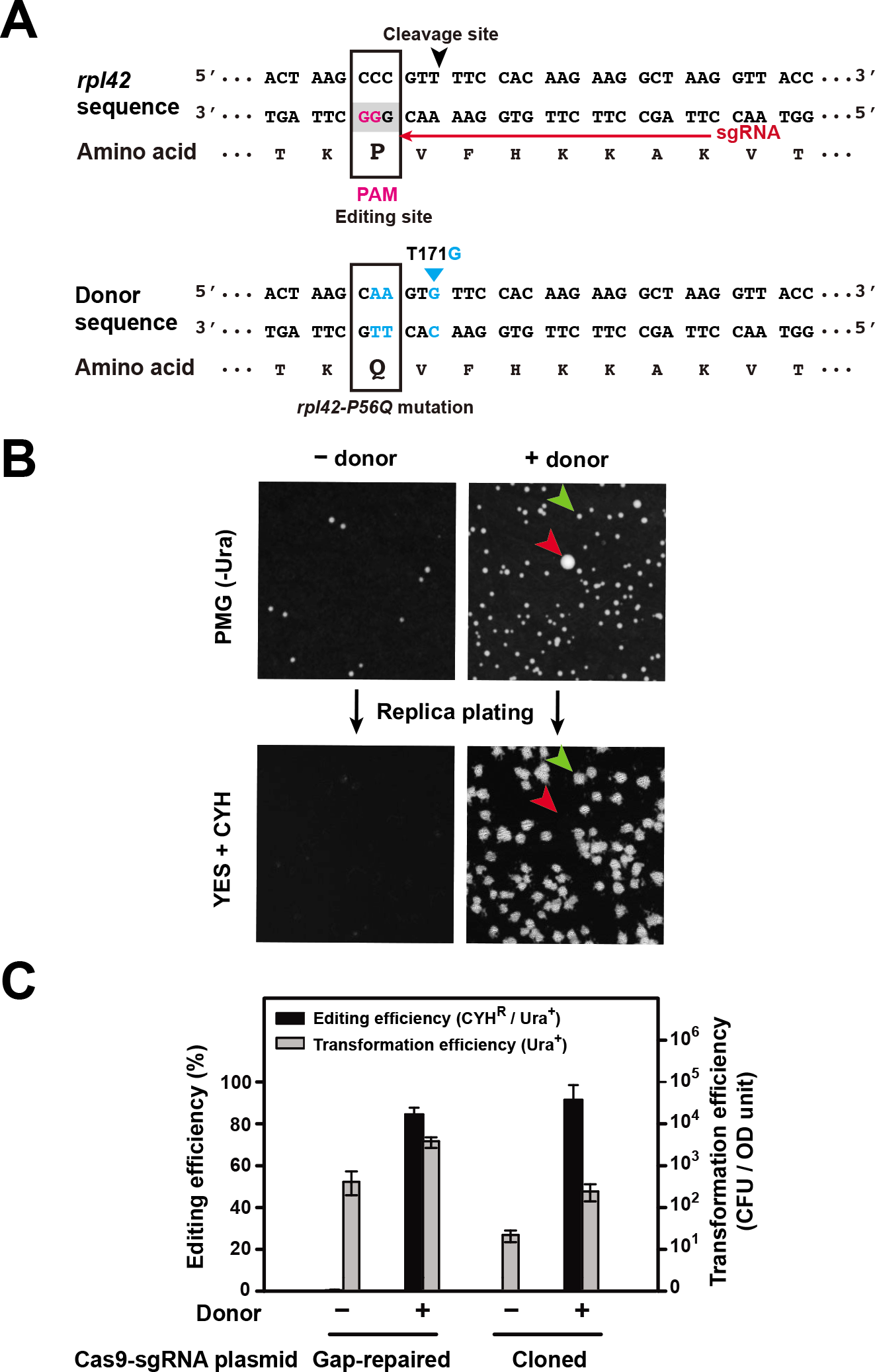
Knock-in of the *rpl42-P56Q* mutation using the split-*ura4* system. A. Sequences of the *rpl42* gene and the donor DNA for introducing the *rpl42-P56Q* mutation. In the *rpl42* sequence, the PAM sequence is shaded in grey and the two invariable guanines in the PAM are highlighted in magenta, the sgRNA target sequence is indicated by a red arrow, and the cleavage site is indicated by a black arrowhead. The editing site (codon to be changed by the knock-in) is boxed. In the donor DNA sequence, the altered nucleotides are highlighted in blue and the altered codon is boxed. T171G is a silent mutation designed to increase the chance of the *rpl42-P56Q* mutation being incorporated during homologous recombination. However, we later found that a donor without the T171G mutation was equally effective in knocking in the *rpl42-P56Q* mutation (data not shown). B. Representative plate images from an *rpl42-P56Q* knock-in experiment using the split-*ura4* system. The images at the top show the Ura^+^ transformant colonies formed on selective plates lacking uracil, and the images at the bottom show the CYH^R^ colonies formed after replica plating transformant colonies onto YES plates containing cycloheximide (CYH, 100 μg/ml). A small transformant colony with the desired genome editing outcome (CYH^R^) is indicated by a green arrowhead, and a rare large transformant colony without the desired genome editing outcome is indicated by a red arrowhead. Only small colonies were considered when calculating the transformation efficiency and the editing efficiency. PCR and Sanger sequencing analysis of 16 CYH^R^ colonies confirmed the presence of the *rpl42-P56Q* mutation in all of their genomes. C. Quantitation of the editing efficiencies and the transformation efficiencies of the *rpl42-P56Q* knock-in experiments. Cas9 and sgRNA were introduced into the cells using either the gap repair procedure (split-*ura4* system, 30 ng of the gapped plasmid and 200 ng of the sgRNA insert) or a cloned Cas9-sgRNA plasmid pDB4284 (30 ng). Donor DNA was provided as two 90-nt complementary oligos (0.3 nmol each). Transformation efficiencies were expressed as colony forming units (CFU) per OD600 unit of cells. Editing efficiencies were expressed as percentages of Ura^+^ transformant colonies that are also CYH^R^ Error bars represent the SD from at least three biological replicates.

We varied the amounts of the gapped plasmid and the sgRNA insert used per transformation (Fig. S1), and found that, taking into account the editing efficiency, the total yield of edited clones, and the reagent cost, 30 ng of the gapped plasmid and 200 ng of the sgRNA insert are an optimal pair of parameters, corresponding to a molar ratio of approximately 1:50, a ratio in line with previous reports on optimal gap repair parameters (Eckert-Boulet et al. 2012; Matsuo et al. 2010). We used such amounts for all ensuing experiments.

Both single-stranded and double-stranded oligos have been used as donor DNA for knocking in mutations (Storici et al. 2003; Chen et al. 2011; DiCarlo et al. 2013). We found that a single 90-nt donor oligo, either the sense oligo or the antisense oligo, can serve as donor to knock in the *rpl42-P56Q* mutation, but knock-in efficiencies were markedly lower than when both oligos were provided together (Fig. S2). A high level of knock-in efficiency was obtained when the two oligos were simply added together into the transformation mix without prior denaturing or annealing (Fig. S2). When varying amounts of donor DNA were tested, we observed that both the editing efficiency and the transformation efficiency increased with the amount of donor DNA up to 0.3 nmol of each oligo, the maximal amount examined (Fig. S3). For economic considerations, we chose to use 0.3 nmol each of two 90-nt complementary oligos for all ensuing knock-in experiments, even though this amount may not be saturating.

#### The split-*bsdMX* system increases the flexibility of the cloning-free procedure

To broaden the utility of the gap-repair based procedure beyond *ura4-D18* strains, we developed another version of it, called the split-*bsdMX* system, which utilizes the blasticidin S resistance selection marker *bsdMX*. In this system, two plasmids pDB4281 and pDB4283 serve as the sources of the gapped plasmid and the template for amplifying the sgRNA insert, respectively (Table 1). Using the *rpl42-P56Q* knock-in again as a proof-of-principle test, we found that the split-*bsdMX* system achieved an editing efficiency comparable to that obtained with the split-*ura4* system, albeit with a notably lower transformation efficiency (Fig. 3).

**Figure 3.**
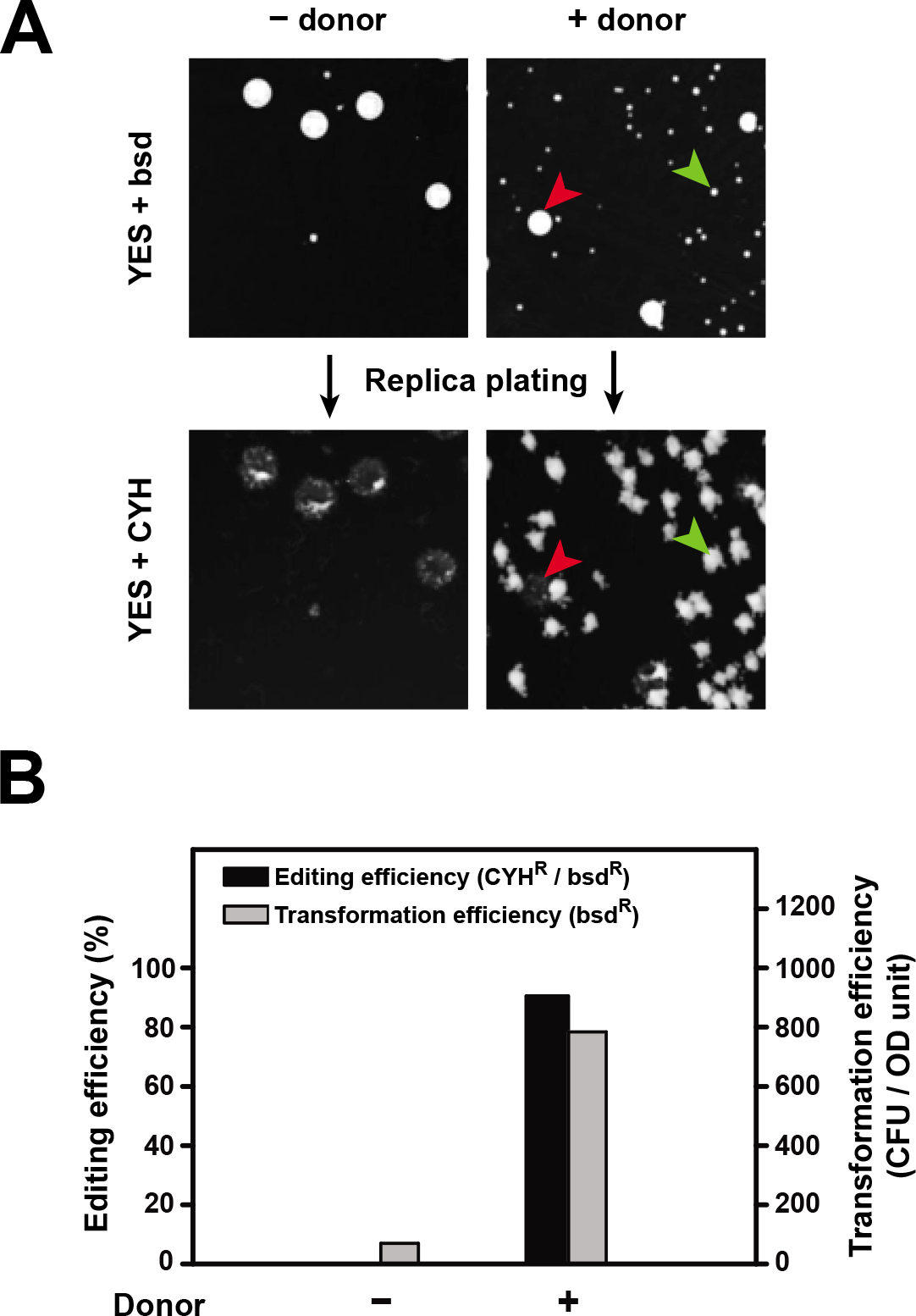
Knock-in of the *rpl42-P56Q* mutation using the split-*bsdMX* system. A. Representative plate images from an *rpl42-P56Q* knock-in experiment using the split-*bsdMX* system. The images at the top show the bsd^R^ transformant colonies formed on YES plates containing blasticidin (bsd, 30 μg/ml), and the images at the bottom show the CYH^R^ colonies formed after replica plating transformant colonies onto YES plates containing CYH (100 μg/ml). A small transformant colony with the desired genome editing outcome (CYH^R^) is indicated by a green arrowhead, and a rare large transformant colony without the desired genome editing outcome is indicated by a red arrowhead. Only small colonies were considered when calculating the transformation efficiency and the editing efficiency. B. Quantitation of the editing efficiencies and the transformation efficiencies of the *rpl42-P56Q* knock-in experiments using the split-*bsdMX* system. 30 ng of the gapped plasmid and 200 ng of the sgRNA insert were used. Donor DNA was provided as two 90-nt complementary oligos (0.3 nmol each).

#### Knock-in of two temperature-sensitive mutations using the cloning-free procedure

To test the versatility and generality of the cloning-free procedure, we applied the split-*ura4* system to knock in two temperature-sensitive (ts) mutations: *tor2-L2048S (tor2-287)* (Hayashi et al. 2007) and *cdc25-C532Y* (*cdc25-22*) (Russell and Nurse 1986; Meyers et al. 2016). Two different sgRNAs were chosen for each mutation (Fig. 4A, 4B, and Table 2). One pair of 90-nt complementary oligos that contains two PAM-disrupting silent mutations was used as the donor DNA for each mutation (Fig. 4A, 4B, and Table S2).

**Figure 4.**
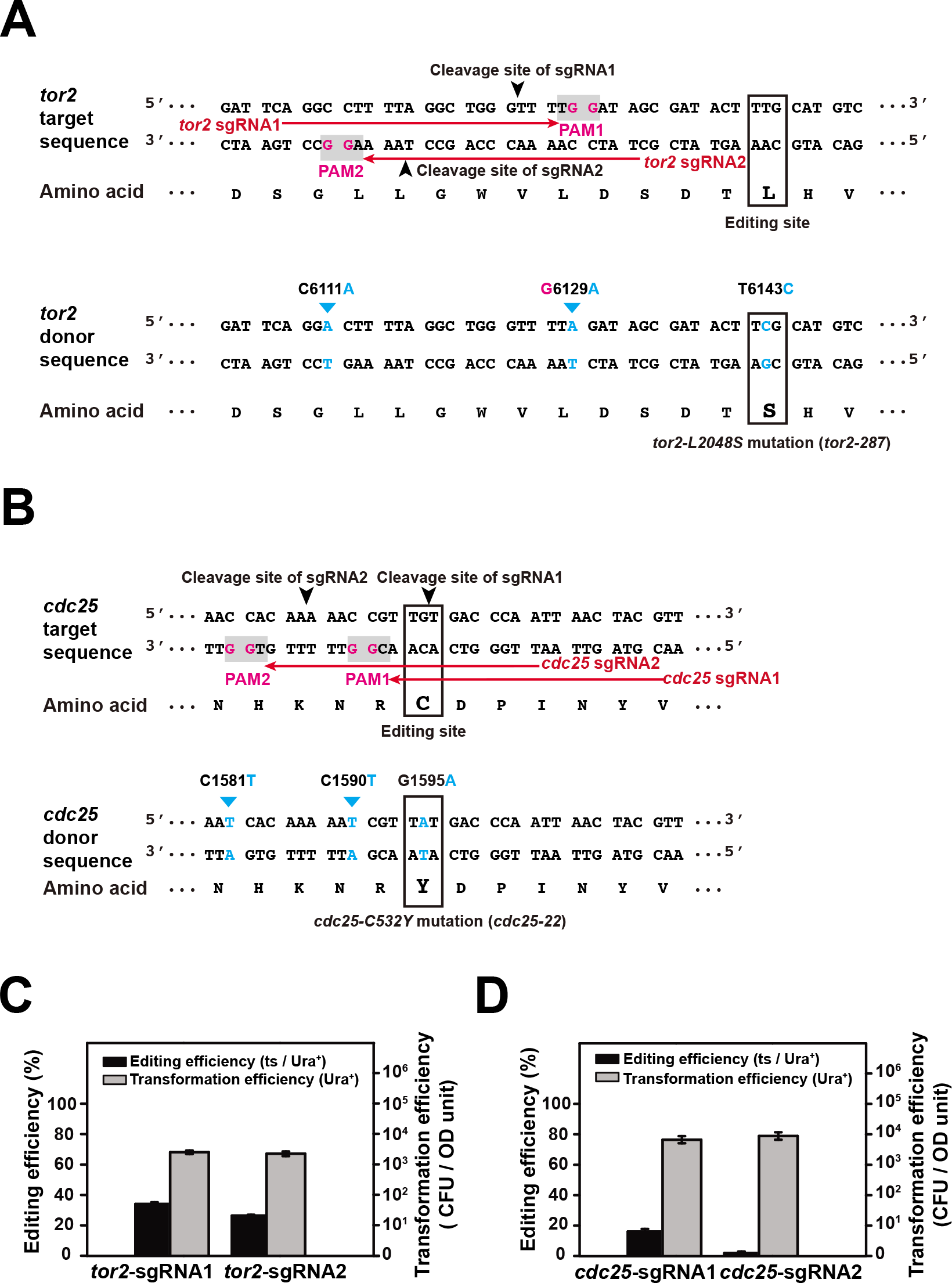
Knock-in of two temperature-sensitive mutations using the split-*ura4* system. A and B. Sequences of the target gene (*tor2* in A and *cdc25* in B) and the donor DNA for introducing a temperature-sensitive (ts) mutation (*tor2-L2048S* in A and *cdc25-C532Y* in B). PAM sequences are shaded in grey; the invariable guanines in the PAMs are highlighted in magenta. The sgRNA target sequences are indicated by red arrows, and the cleavage sites are indicated by black arrowheads. The editing site (codon to be changed by the knock-in) is boxed. In the donor DNA sequence, the altered nucleotides are highlighted in blue and the altered codon is boxed. C6111A and G6129A in *tor2* and C1581T and C1590T in *cdc25* are cleavage-blocking silent mutations that disrupt the PAMs but do not change the amino acid sequences. C. Quantitation of the editing efficiencies and the transformation efficiencies of the *tor2-L2048S* knock-in. PCR and Sanger sequencing analysis of 11 temperature-sensitive colonies confirmed the presence of the *tor2-L2048S* mutation in all of their genomes. D. Quantitation of the editing efficiencies and the transformation efficiencies of the *cdc25-C532Y* knock-in. PCR and Sanger sequencing analysis of 10 temperature-sensitive colonies confirmed the presence of the *cdc25-C532Y* mutation in all of their genomes.

Compared to the *rpl42-P56Q* knock-in, we obtained more modest levels of knock-in efficiencies (34% and 27% respectively for the two sgRNAs) when knocking in the *tor2-L2048S* mutation (Fig. 4C). We suspected that the lower efficiencies may be due to the cleavage sites being farther away from the editing site (18 bp and 26 bp respectively for the two *tor2* sgRNAs versus 3 bp for the *rpl42* sgRNA). Indeed, when a number of randomly selected transformants without the ts phenotype were examined by PCR and Sanger sequencing analysis, we found that nearly all them (11/11 for sgRNA1 and 7/8 for sgRNA2) harbored the PAM-disrupting silent mutation (Table S3), indicating that Cas9-mediated cleavage and donor recombination were highly efficient but the incorporation of distal mutations only occurred in a fraction of the recombination products.

Knock-in of the *cdc25-C532Y* mutation using sgRNA1 only achieved a moderate editing efficiency (16%), despite the cleavage site being immediately adjacent to the editing site, suggesting that the cleavage may be inefficient (Fig. 4D). Consistent with this possibility, all 8 randomly chosen non-ts transformant clones had the wild-type *cdc25* sequence (Table S3). Even worse editing efficiency (2%) was obtained using sgRNA2, which induces cleavage 8 bp away from the editing site (Fig. 4D). In this case, cleavage efficiency did not appear to be very poor, as approximately 40% (5/13) of the transformants without the ts phenotype contained the PAM-disrupting mutation (Table S3).

Besides the temperature sensitivity, the *tor2-L2048S* and *cdc25-C532Y* mutants obtained by Cas9-mediated knock-in exhibited the other expected phenotypes, including rapamycin sensitivity and short-cell morphology for the former, and cell elongation phenotype for the latter (Figure S4 and S5).

These experiments demonstrated that the cloning-free procedure is universally applicable for point mutation knock-in. Because sgRNA cleavage efficiency is variable, using more than one sgRNA is a worthwhile strategy to ensure knock-in success. Close proximity between the cleavage site and the editing site is beneficial, but not absolutely required for the Cas9-mediated knock-in of point mutations.

#### N-terminal tagging using the cloning-free procedure

We next tested if this gap-repair-based CRISPR/Cas9 system could be used for N-terminal tagging. We chose to add an mECtrine fluorescent protein tag to Ypt7, a protein functioning in membrane fusion events involving vacuoles. Ypt7 is post-translationally modified at the C-terminus by prenylation and cannot tolerate C-terminal tagging (Landgraf et al. 2016). We selected two different sgRNA target sequences, both of which are disrupted if tagging is successful (Fig. 5A). Around 60% tagging efficiencies were achieved using sgRNA1 together with donor PCR products containing homology arms with lengths ranging from 50 to 200 bp (Fig. 5B). In contrast, sgRNA2 completely failed to generate correctly tagged clones, reinforcing the notion that sgRNAs can be variable in their targeting efficiency. mECtrine-tagged Ypt7 obtained by Cas9-mediated tagging exhibited the expected vacuole localization when observed by live cell imaging (Fig. S6) (Kashiwazaki et al. 2005).

**Figure 5.**
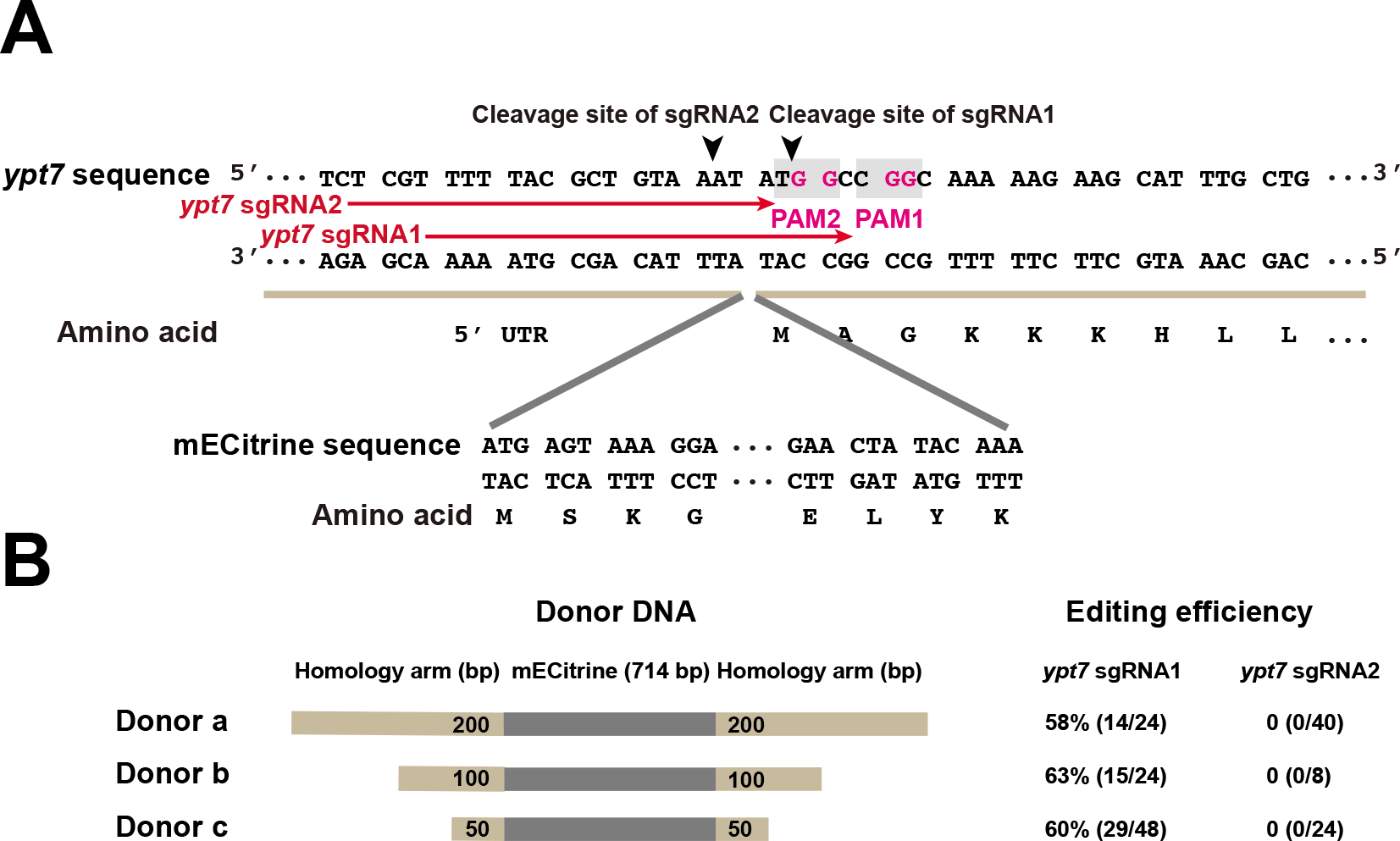
Tagging the N-terminus of Ypt7 using the split-*ura4* system. A. Sequences of the N-terminal region of the *ypt7* gene and the mECtrine tag. PAM sequences are shaded in grey; the invariable guanines in the PAMs are highlighted in magenta. The sgRNA target sequences are indicated by red arrows, and the cleavage sites are indicated by black arrowheads. B. Schematic of donor DNAs (not to scale) and their editing efficiencies when used with two different sgRNAs. Donor DNAs are composed of the mECtrine tag (shown in grey) and flanking homology arms (shown in light brown). The lengths of the homology arms are 200 bp, 100 bp, and 50 bp in donor a, b, and c, respectively. Editing efficiencies were assessed by live cell imaging analysis of individual Ura^+^ transformant clones. Correct editing resulted in the vacuole membrane being labeled by mECtrine (Figure S6).

#### Genomic sequence deletion using the cloning-free procedure

Lastly, we tested whether the gap-repair based system can be used in genomic sequence deletion. We used the split-*bsdMX* system to generate the *mat1*-Δ*17* deletion, which removes a 140-bp sequence from the *mat1* locus and converts homothallic strains that can efficiently switch mating type to heterothallic strains that cannot switch mating type (Arcangioli and Klar 1991). Two sgRNA target sequences were chosen within the 140-bp sequence (Fig. 6A and Table 2). The donor DNA was a pair of 90-nt complementary oligos that each consists of two 45-nt homology arms flanking the sequence to be deleted (Fig. 6A). Conversion to heterothallism was scored as bsd^R^ transformants that when replica-plated on a sporulating plate, cannot be darkly stained by iodine vapor (Fig. 6B) (Forsburg and Rhind 2006). Greater than 90% conversion efficiencies were achieved on JB938, a natural isolate of *S. pombe* (Fig. 6B and 6C), demonstrating the utility of the split-*bsdMX* system in editing the genomes of non-laboratory *S. pombe* strains, which have become increasingly useful for genetic and evolutionary studies (Jeffares et al. 2015; Hu et al. 2015).

**Figure 6.**
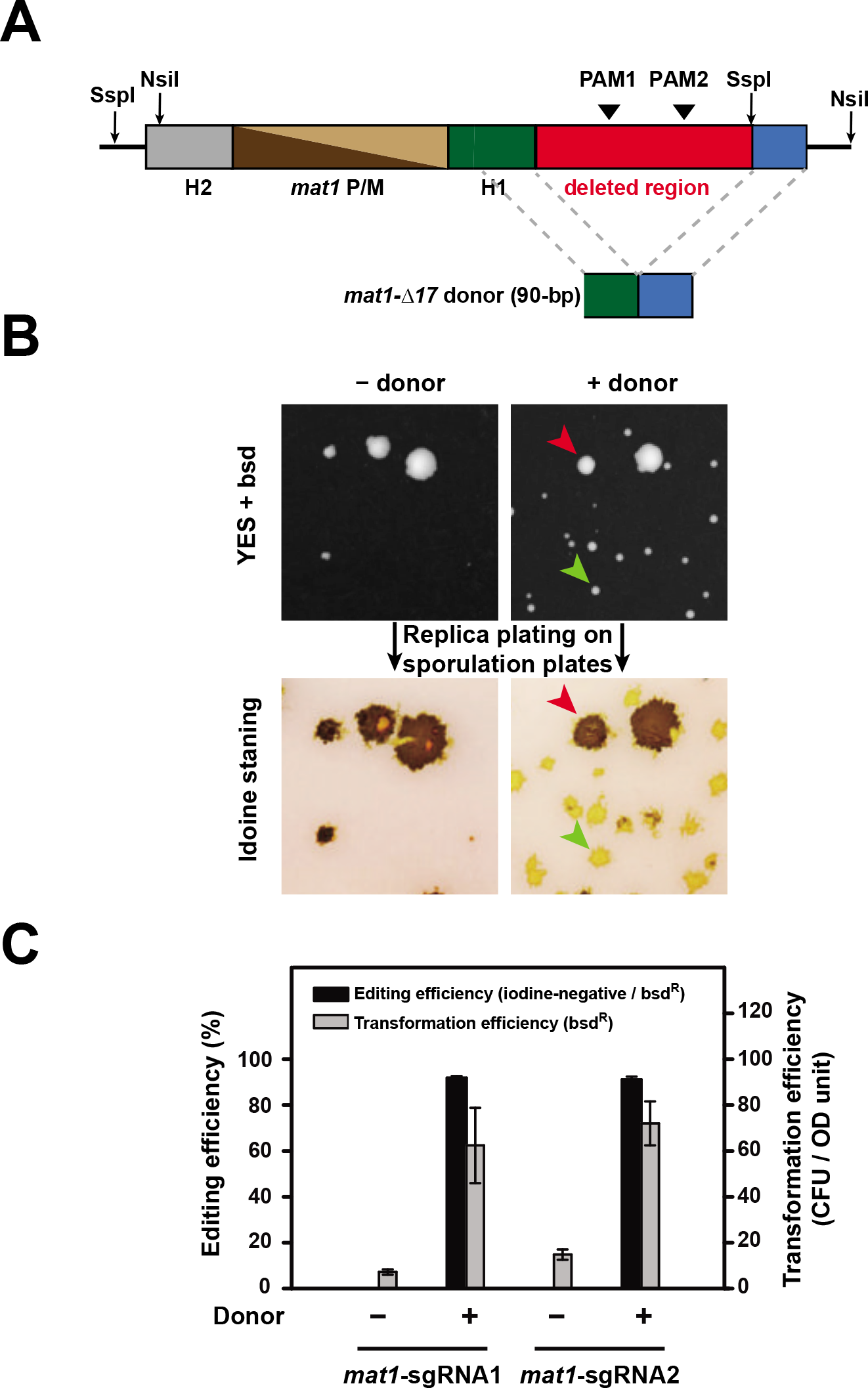
Use of the split-*bsdMX* system in genomic sequence deletion. A. Schematic of the *mat1* locus of a homothallic strain and the donor for generating the *mat1-A17* deletion (not to scale). Two PAM sites were chosen within the to-be-deleted 140-bp sequence, which is situated between the H1 homology box and an SspI restriction site. Two 90-nt complementary oligos each composed of two 45-nt sequences flanking the 140-bp sequence were used as the donor. B. Representative plate images from a *mat1*-Δ*17* deletion experiment using the split-*bsdMX* system. The parental strain is JB938, a homothallic *S. pombe* natural isolate (Jeffares et al. 2015). The images at the top show the bsd^R^ transformant colonies formed on YES plates containing blasticidin (bsd, 30 μg/ml), and the images at the bottom show the iodine staining of transformant colonies replica-plated on sporulation plates (SPAS). A small transformant colony with the desired genome editing outcome (iodine-negative, i.e., cannot be darkly stained by iodine vapor) is indicated by a green arrowhead, and a rare large transformant colony without the desired genome editing outcome is indicated by a red arrowhead. Only small colonies were considered when calculating the transformation efficiencies and the editing efficiencies. C. Quantitation of the editing efficiencies and the transformation efficiencies of the *mat1-A17* deletion in JB938. PCR and Sanger sequencing analysis of 8 iodine-negative colonies confirmed the presence of the *mat1*-Δ*17* deletion in their genomes.

### DISCUSSION

Here, we developed a cloning-free CRISPR/Cas9 method based on gap repair in fission yeast cells. Previously, gap-repair-based CRISPR/Cas9 procedures have been developed in budding yeast to achieve cloning-free genome editing (Horwitz et al. 2015; Mans et al. 2015). The gapped plasmids used in those studies contain intact selection markers and thus, if recircularize through NHEJ-mediated DSB repair, can transform budding yeast cells without incorporating the sgRNA insert. Indeed, one of the studies reported a high background transformation efficiency when the sgRNA insert was omitted (Mans et al. 2015). In theory, this problem can be mitigated in budding yeast by using gapped plasmids with blunt ends, as the budding yeast NHEJ machinery cannot efficiently recircularize linearized plasmids bearing blunt ends (Boulton and Jackson 1996). However, such a remedy is not applicable to fission yeast, because gapped plasmids with either cohesive or blunt ends, if containing intact selection markers, can transform fission yeast cells as efficiently as circular plasmids (Goedecke et al. 1994; Wilson et al. 1999; Manolis et al. 2001). The split-marker design we implemented in this study successfully circumvents this background transformation issue and should be useful to other organisms where NHEJ-mediated plasmid recircularization can compete with gap repair.

It was reported that, in budding yeast, the selection of a gap-repair outcome during transformation enhanced the efficiency of recombination-mediated genome editing, possibly due to an enrichment of cells in S/G2/M phase of the cell cycle, which have a stronger homologous recombination activity than cells in the G1 phase of the cell cycle (Horwitz et al. 2015). We do not know whether such an enhancement effect occurs in fission yeast, because in the only head-to-head comparison between the gap-repair-based method and the circular-plasmid-based method (Figure 2), both methods achieved near saturating levels of editing efficiency. We suspect that such an effect, if exists, may not be as pronounced as in budding yeast, because fission yeast has a much shorter G1 phase than budding yeast and most of the fission yeast cells in an asynchronous culture are in the G2 phase. On the other hand, we did uncover an unexpected advantage of the gap-repair-based method, as it yielded a more than 10 times higher transformation efficiency compared to the circular-plasmid-based method (Figure 2). We speculate that the delay of sgRNA (and perhaps also Cas9) expression in the gap-repair procedure, due to the time needed to complete gap repair, may allow the cells to first recover from the stress of the transformation procedure before encountering the stresses generated by Cas9 and sgRNA.

One practical issue of using Cas9 in fission yeast is its sgRNA-independent growth-inhibition effect, which results in a longer-than-usual incubation time for transformant colonies to form (6-8 days with the split-*ura4* system). The sgRNA-independent cytotoxicity of Cas9 has been reported in a number of organisms, including the bacteria *Clostridium pasteurianum* (Pyne et al. 2016) and *Clostridium acetobutylicum* (Bruder et al. 2016), the cyanobacterium *Synechococcus elongatus* (Wendt et al. 2016), the budding yeast *Saccharomyces cerevisiae* (Ryan et al. 2014; Ronda et al. 2015; Generoso et al. 2016), the fission yeast *S. pombe* (Jacobs et al. 2014), and the single-cell alga *Chlamydomonas reinhardtii* (Jiang et al. 2014). In the cases of *S. pombe* and *Chlamydomonas reinhardtii*, it has been shown that the cytotoxicity is independent of the Cas9 nuclease activity (Jiang et al. 2014; Ciccaglione 2015). We have attempted to circumvent this issue by using weaker promoters or repressible promoters to drive the expression of Cas9, but have thus far failed to identify a condition where reduced cytotoxicity is not accompanied by a severe reduction of editing efficiency (data not shown). Alternative strategies, such as appending an inducible degron to Cas9 (Senturk et al. 2017), may need to be explored in the future to further improve the application of CRISPR/Cas9-based genome editing in *S. pombe*.

## ACKNOWLEDGMENTS

We thank Mikel Zaratiegui for making pMZ374 and other Cas9-related plasmids available through Addgene, Jürg Bähler for making the natural isolate strains available through YGRC/NBRP, and YGRC/NBRP for providing the JB938 strain. This work was supported by grants from the Ministry of Science and Technology of the People’s Republic of China and the Beijing municipal government.

## SUPPLEMENTARY MATERIALS

Figures S1–S6, Tables S1–S3, and Supplementary Data File 1-5 in the supplementary file.

## References

Aoi, Y., M. Sato, T. Sutani, K. Shirahige, T.M. Kapoor et al., 2014 Dissecting the first and the second meiotic divisions using a marker-less drug-hypersensitive fission yeast. Cell Cycle 13 (8):1327-1334.

Arcangioli, B., and A.J. Klar, 1991 A novel switch-activating site (SAS1) and its cognate binding factor (SAP1) required for efficient mat1 switching in Schizosaccharomyces pombe. Embo J 10 (10):3025-3032.

Boulton, S.J., and S.P. Jackson, 1996 Identification of a Saccharomyces cerevisiae Ku80 homologue: roles in DNA double strand break rejoining and in telomeric maintenance. Nucleic Acids Res 24 (23):4639-4648.

Bruder, M.R., M.E. Pyne, M. Moo-Young, D.A. Chung, and C.P. Chou, 2016 Extending CRISPR-Cas9 Technology from Genome Editing to Transcriptional Engineering in the Genus Clostridium. Appl Environ Microbiol 82 (20):6109-6119.

Chen, F., S.M. Pruett-Miller, Y. Huang, M. Gjoka, K. Duda et al., 2011 High-frequency genome editing using ssDNA oligonucleotides with zinc-finger nucleases. Nat Methods 8 (9):753-755.

Chino, A., K. Watanabe, and H. Moriya, 2010 Plasmid construction using recombination activity in the fission yeast Schizosaccharomyces pombe. PLoS One 5 (3):e9652.

Ciccaglione, K.M., 2015 Adaptation of CRISPR/Cas9 to improve the experimental utility of the model system schizosaccharomyces pombe. Master’s thesis Rutgers University.

Colon, M., and N.C. Walworth, 2004 Use of in vivo gap repair for isolation of mutant alleles of a checkpoint gene. Methods Mol Biol 241:175-187.

DiCarlo, J.E., J.E. Norville, P. Mali, X. Rios, J. Aach et al., 2013 Genome engineering in Saccharomyces cerevisiae using CRISPR-Cas systems. Nucleic Acids Res 41 (7):4336-4343.

Doench, J.G., N. Fusi, M. Sullender, M. Hegde, E.W. Vaimberg et al., 2016 Optimized sgRNA design to maximize activity and minimize off-target effects of CRISPR-Cas9. Nat Biotechnol 34 (2):184-191.

Doudna, J.A., and E. Charpentier, 2014 Genome editing. The new frontier of genome engineering with CRISPR-Cas9. Science 346 (6213):1258096.

Eckert-Boulet, N., M.L. Pedersen, B.O. Krogh, and M. Lisby, 2012 Optimization of ordered plasmid assembly by gap repair in Saccharomyces cerevisiae. Yeast 29 (8):323-334.

Fennessy, D., A. Grallert, A. Krapp, A. Cokoja, A.J. Bridge et al., 2014 Extending the Schizosaccharomyces pombe molecular genetic toolbox. PLoS One 9 (5):e97683.

Fernandez, R., and J. Berro, 2016 Use of a fluoride channel as a new selection marker for fission yeast plasmids and application to fast genome editing with CRISPR/Cas9. Yeast 33 (10):549-557.

Forsburg, S.L., and N. Rhind, 2006 Basic methods for fission yeast. Yeast 23 (3):173-183.

Gao, Y., and Y. Zhao, 2014 Self-processing of ribozyme-flanked RNAs into guide RNAs in vitro and in vivo for CRISPR-mediated genome editing. J Integr Plant Biol 56 (4): 343-349.

Generoso, W.C., M. Gottardi, M. Oreb, and E. Boles, 2016 Simplified CRISPR-Cas genome editing for Saccharomyces cerevisiae. J Microbiol Methods 127:203-205.

Gietz, R.D., and R.A. Woods, 2006 Yeast transformation by the LiAc/SS Carrier DNA/PEG method. Methods Mol Biol 313:107-120.

Goedecke, W., P. Pfeiffer, and W. Vielmetter, 1994 Nonhomologous DNA end joining in Schizosaccharomyces pombe efficiently eliminates DNA double-strand-breaks from haploid sequences. Nucleic Acids Res 22 (11):2094-2101.

Grimm, C., J. Kohli, J. Murray, and K. Maundrell, 1988 Genetic engineering of Schizosaccharomyces pombe: a system for gene disruption and replacement using the ura4 gene as a selectable marker. Mol Gen Genet 215 (1):81-86.

Hayashi, T., M. Hatanaka, K. Nagao, Y. Nakaseko, J. Kanoh et al., 2007 Rapamycin sensitivity of the Schizosaccharomyces pombe tor2 mutant and organization of two highly phosphorylated TOR complexes by specific and common subunits. Genes Cells 12 (12):1357-1370.

Horwitz, A.A., J.M. Walter, M.G. Schubert, S.H. Kung, K. Hawkins et al., 2015 Efficient Multiplexed Integration of Synergistic Alleles and Metabolic Pathways in Yeasts via CRISPR-Cas. Cell Syst 1 (1):88-96.

Hu, W., F. Suo, and L.L. Du, 2015 Bulk Segregant Analysis Reveals the Genetic Basis of a Natural Trait Variation in Fission Yeast. Genome Biol Evol 7 (12):3496-3510.

Jacobs, J.Z., K.M. Ciccaglione, V. Tournier, and M. Zaratiegui, 2014 Implementation of the CRISPR-Cas9 system in fission yeast. Nat Commun 5:5344.

Jeffares, D.C., C. Rallis, A. Rieux, D. Speed, M. Prevorovsky et al., 2015 The genomic and phenotypic diversity of Schizosaccharomyces pombe. Nat Genet 47 (3):235-241.

Jiang, W., A.J. Brueggeman, K.M. Horken, T.M. Plucinak, and D.P. Weeks, 2014 Successful transient expression of Cas9 and single guide RNA genes in Chlamydomonas reinhardtii. Eukaryot Cell 13 (11):1465-1469.

Jinek, M., K. Chylinski, I. Fonfara, M. Hauer, J.A. Doudna et al., 2012 A programmable dual-RNA-guided DNA endonuclease in adaptive bacterial immunity. Science 337 (6096):816-821.

Kashiwazaki, J., T. Nakamura, T. Iwaki, K. Takegawa, and C. Shimoda, 2005 A role for fission yeast Rab GTPase Ypt7p in sporulation. Cell Struct Funct 30 (2):43-49.

Kelly, D.A., and C.S. Hoffman, 2002 Gap repair transformation in fission yeast to exchange plasmid-selectable markers. Biotechniques 33 (5):978, 980, 982.

Kostrub, C.F., E.P. Lei, and T. Enoch, 1998 Use of gap repair in fission yeast to obtain novel alleles of specific genes. Nucleic Acids Res 26 (20):4783-4784.

Landgraf, D., D. Huh, E. Hallacli, and S. Lindquist, 2016 Scarless Gene Tagging with One-Step Transformation and Two-Step Selection in Saccharomyces cerevisiae and Schizosaccharomyces pombe. PLoS One 11 (10):e0163950.

Mali, P., L. Yang, K.M. Esvelt, J. Aach, M. Guell et al., 2013 RNA-guided human genome engineering via Cas9. Science 339 (6121):823-826.

Manolis, K.G., E.R. Nimmo, E. Hartsuiker, A.M. Carr, P.A. Jeggo et al., 2001 Novel functional requirements for non-homologous DNA end joining in Schizosaccharomyces pombe. Embo J 20 (1-2):210-221.

Mans, R., H.M. van Rossum, M. Wijsman, A. Backx, N.G. Kuijpers et al., 2015 CRISPR/Cas9: a molecular Swiss army knife for simultaneous introduction of multiple genetic modifications in Saccharomyces cerevisiae. FEMS Yeast Res 15 (2).

Mao, Y., J. Lin, A. Zhou, K. Ji, J.S. Downey et al., 2011 Quikgene: a gene synthesis method integrated with ligation-free cloning. Anal Biochem 415 (1):21-26.

Matsuo, Y., H. Kishimoto, T. Horiuchi, K. Tanae, and M. Kawamukai, 2010 Simple and effective gap-repair cloning using short tracts of flanking homology in fission yeast. Biosci Biotechnol Biochem 74 (3):685-689.

Meyers, A., Z.P. Del Rio, R.A. Beaver, R.M. Morris, T.M. Weiskittel et al., 2016 Lipid Droplets Form from Distinct Regions of the Cell in the Fission Yeast Schizosaccharomyces pombe. Traffic 17 (6):657-669.

Pyne, M.E., M.R. Bruder, M. Moo-Young, D.A. Chung, and C.P. Chou, 2016 Harnessing heterologous and endogenous CRISPR-Cas machineries for efficient markerless genome editing in Clostridium. Sci Rep 6:25666.

Rodriguez-Lopez, M., C. Cotobal, O. Fernandez-Sanchez, N. Borbaran Bravo, R. Oktriani et al., 2016 A CRISPR/Cas9-based method and primer design tool for seamless genome editing in fission yeast. Wellcome Open Res 1:19.

Roguev, A., M. Wiren, J.S. Weissman, and N.J. Krogan, 2007 High-throughput genetic interaction mapping in the fission yeast Schizosaccharomyces pombe. Nat Methods 4 (10):861-866.

Ronda, C., J. Maury, T. Jakociunas, S.A. Jacobsen, S.M. Germann et al., 2015 CrEdit: CRISPR mediated multi-loci gene integration in Saccharomyces cerevisiae. Microb Cell Fact 14:97.

Russell, P., and P. Nurse, 1986 cdc25+ functions as an inducer in the mitotic control of fission yeast. Cell 45 (1):145-153.

Ryan, O.W., J.M. Skerker, M.J. Maurer, X. Li, J.C. Tsai et al., 2014 Selection of chromosomal DNA libraries using a multiplex CRISPR system. Elife 3.

Sander, J.D., and J.K. Joung, 2014 CRISPR-Cas systems for editing, regulating and targeting genomes. Nat Biotechnol 32 (4):347-355.

Senturk, S., N.H. Shirole, D.G. Nowak, V. Corbo, D. Pal et al., 2017 Rapid and tunable method to temporally control gene editing based on conditional Cas9 stabilization. Nat Commun 8:14370.

Shirai, A., M. Sadaie, K. Shinmyozu, and J. Nakayama, 2010 Methylation of ribosomal protein L42 regulates ribosomal function and stress-adapted cell growth. J Biol Chem 285 (29):22448-22460.

Storici, F., C.L. Durham, D.A. Gordenin, and M.A. Resnick, 2003 Chromosomal site-specific double-strand breaks are efficiently targeted for repair by oligonucleotides in yeast. Proc Natl Acad Sci U S A 100 (25):14994-14999.

Wang, H., M. La Russa, and L.S. Qi, 2016 CRISPR/Cas9 in Genome Editing and Beyond. Annu Rev Biochem 85:227-264.

Wendt, K.E., J. Ungerer, R.E. Cobb, H. Zhao, and H.B. Pakrasi, 2016 CRISPR/Cas9 mediated targeted mutagenesis of the fast growing cyanobacterium Synechococcus elongatus UTEX 2973. Microb Cell Fact 15 (1):115.

Wilson, S., N. Warr, D.L. Taylor, and F.Z. Watts, 1999 The role of Schizosaccharomyces pombe Rad32, the Mre11 homologue, and other DNA damage response proteins in non-homologous end joining and telomere length maintenance. Nucleic Acids Res 27 (13):2655-2661.

